# Parallel valence processing alterations associated with compulsive behavior in SAPAP3 knockout mice and human OCD

**DOI:** 10.1101/2021.06.04.447162

**Authors:** Bridget L. Kajs, Peter J. van Roessel, Gwynne L. Davis, Leanne M. Williams, Carolyn I. Rodriguez, Lisa A. Gunaydin

## Abstract

Abnormalities in valence processing – the processing of aversive or appetitive stimuli – may be an underrecognized component of obsessive-compulsive disorder (OCD). Independent experimental paradigms have suggested disturbance of emotional valence systems in OCD, yet no standardized assay has been employed to assess both negative and positive valence processing in clinical studies of OCD patients, either at baseline or in response to therapeutic interventions. Additionally, preclinical rodent models are critical for treatment discovery in OCD, yet investigations examining whether rodent models of compulsive behavior similarly show alterations in valence systems have been limited. We sought to establish paradigms for assessing valence processing across both human OCD patients and in a preclinical rodent model: in OCD patients, we used validated behavioral tests to assess explicit and implicit processing of fear-related facial expressions (negative valence) and socially-rewarding happy expressions (positive valence); in the SAPAP3 knockout (KO) mouse model of compulsive behavior, we used auditory fear conditioning and extinction (negative valence) and reward-based operant conditioning (positive valence). We find that OCD patients show enhanced negative and impaired positive valence processing, and that performance on valence processing tasks correlates with clinical measures of OCD severity. We further find that SAPAP3 KO mice show heightened negative and impaired positive valence processing alterations similar to those of OCD patients. Our results show parallel valence processing abnormalities in OCD patients and a preclinical rodent model of compulsive behavior, and suggest valence processing alterations as novel therapeutic targets across a translational research spectrum.

## INTRODUCTION

Neural systems that process valence (i.e., positive and negative affect) are crucial for orchestrating adaptive behavioral responses to individuals, objects, or events with emotional salience. Dysfunction or imbalance of valence processing systems contributes to many psychiatric disorders, for example the overactive threat appraisal of anxiety disorders or the overactive reward salience of addictive disorders. Obsessive-compulsive disorder (OCD), a common and disabling disorder characterized by intrusive thoughts and repetitive behaviors ^1^, is one psychiatric condition in which valence processing abnormalities may play a critical role in disease pathology. However, the relationship between valence processing abnormalities and compulsive behavior is poorly understood.

Despite the establishment of negative valence systems and positive valence systems as NIMH RDoC research domains ^2^, few translational studies have explicitly assessed valence processing in OCD. Nonetheless, many lines of research suggest that alterations of both negative and positive valence processing may be characteristic of the disorder. Individuals with OCD overestimate threat ^3^, endorse greater intolerance of uncertainty ^4,5^, and are impaired in their ability to distinguish between safe and threatening stimuli ^6^. OCD patients also show an elevated startle response ^7^, hyperactivation of fear circuitry in response to OCD-relevant stimuli ^8^, and impaired fear extinction learning ^9–12^, further suggesting overactive threat processing as a negative valence alteration. With regard to positive valence systems, individuals with OCD exhibit greater levels of anhedonia, suggesting deficits in reward valuation ^13–16^. Additionally, OCD patients show abnormalities in responsiveness during reward anticipation ^17^ and impairments in their ability to process reward feedback ^18,19^.

Valence processing abnormalities may relate in fundamental ways to the behaviors characteristic of OCD. Overactive threat detection may drive behavioral avoidance and compulsions aimed at mitigating threat or decreasing anxiety ^4,20^, while deficits in reward processing may impair action-outcome contingency learning ^18,19,21^ and bias OCD patients towards habitual over goal-directed behavior ^22–25^. Furthermore, the proneness to habitual behavior observed in OCD may further be driven by hyperactive negative valence processing ^26,27^.

Measurable alterations of both positive and negative valence processing may have significance as markers of illness severity, targets for therapeutic intervention, or as predictors of treatment response. Indeed, exposure with response prevention (ERP), the first-line and most effective behavioral therapy for OCD, targets negative valence abnormalities in OCD through the extinction of excessive threat processing over time ^28^, and excessive fear behavior has been observed to predict worse treatment outcomes with selective serotonin reuptake inhibitors (SSRIs) or cognitive behavioral therapy (CBT) ^12,29^. Nonetheless, measures of valence processing have not typically been incorporated in clinical trials of OCD therapeutics, due to the lack of standardized, easily reproducible means for assessing both positive and negative valence processing in clinical populations.

Studies examining valence processing alterations in model systems relevant to OCD are even more limited. Mouse strains in which specific genetic deletions lead to compulsive-like repetitive behaviors have advanced pathophysiological understanding of OCD ^30–33^. In one of these well established models, the SAPAP3 knockout (KO) mouse, loss of *Sapap3* gene expression leads to impaired frontostriatal synaptic transmission and excessive and detrimental grooming^34^. OCD pathology has similarly been tied to dysregulated frontostriatal circuits ^35–37^ and treatments that improve OCD symptoms reduce abnormal frontostriatal functional connectivity^38^. These behavioral and physiological similarities between the SAPAP3 KO model and OCD suggest that understanding valence processing in SAPAP3 mice could further advance translational science relevant to OCD.

In this study, we sought to examine valence processing across both OCD patients and the SAPAP3 KO mouse model. In OCD patients, we utilized a standardized, computer-based cognitive testing platform to examine both explicit (conscious) and implicit (unconscious) processing of emotional face stimuli. We specifically assessed responses to faces expressing fear and happiness, given evidence that they potently engage brain regions involved in the processing of threat (negative valence) and reward (positive valence), respectively ^39^. In SAPAP3 KO mice, we examined alterations in valence processing using a classical auditory fear conditioning and extinction paradigm (negative valence) as well as a reward-based operant conditioning paradigm (positive valence). We find that OCD patients and SAPAP3 KO mice show consistent trends towards enhanced negative valence processing and impaired positive valence processing. These results confirm the translational potential of our valence processing assays and suggest future biological investigation utilizing these assays is warranted.

## METHODS

### Human subjects

Stanford University Administrative Panel for the Protection of Human Subjects (Institutional Review Board) approval was obtained for all procedures and all participants gave informed consent. Participants ages 18 to 65 seeking to participate in clinical treatment studies of OCD were recruited through local, radio, and internet advertisement between October 2016 and October 2019. Participants were required to have OCD as assessed by the structured clinical interview for DSM-5 ^40^ with at least moderate symptoms (Yale-Brown Obsessive-Compulsive Scale (Y-BOCS) ^41^ score > 16 and recurrent intrusions > 8 hours daily) and to have been free of psychotropic medication for 12 weeks. All diagnostic or clinician-rated assessments were performed by trained PhD or MD-level assessors. Participants were excluded if they had primary hoarding/saving obsessions, severe depression, bipolar disorder, history of psychosis, seizure disorder, prior head injury or other neurological disorders, current moderate or history of severe substance use disorder, or severe medical comorbidity.

### Animals

SAPAP3 wild-type (WT) and knock-out (KO) mice (B6.129-*Dlgap3tm1Gfng*/J) bred on a C57/BL6 background were used for all experiments. Genotypes of all animals were confirmed using PCR amplification. Animals were raised in normal light conditions (12:12 light/dark cycle), and given food and water ad libitum. All experiments were conducted in accordance with procedures established by the Institutional Animal Care and Use Committee at the University of California, San Francisco. Experimenters were blinded to genotypes during all behavioral assays.

### Clinical Measures

Depression Anxiety Stress Scales (DASS). The DASS ^42^ is a 21-item self-report measure assessing depression, anxiety, and stress over the past week. Scores range from 0 to 42 for each of three subscales. DASS subscales show good psychometric properties in clinical and non-clinical cohorts ^43^.

Yale Brown Obsessive-Compulsive Scale (Y-BOCS). The Y-BOCS is a clinician-rated instrument that scores both obsessions (5 items) and compulsions (5 items) for a total OCD severity score ranging between 0 and 40. Obsessions and Compulsions subscales range from 0-20. The Y-BOCS is well-validated with good psychometric properties and is the most widely used rating scale of OCD severity^41^.

### Emotion processing tasks

Data used to quantify human emotion processing were obtained using a computer-based testing battery (‘WebNeuro’) that allows standardized assessment of behavioral performance on both emotional and general cognitive tasks ^44^. WebNeuro tasks have been validated against gold-standard neuropsychological tests assessing equivalent constructs and demonstrate sound psychometric properties ^44,45^. Subject-level performance is quantified as a z-score with reference to age-, biological sex-, and years of education-matched norms derived from a healthy norm cohort of n=1317 ^46,47^. WebNeuro tests are completed in a single, undistracted sitting of approximately 45 minutes. Participants were offered the opportunity to complete testing either at home or in a clinic office.

WebNeuro tasks used in this study included i) an *explicit emotion processing* task, assessing participants’ conscious verbal labeling of emotional face stimuli, and, ii) a subsequent *implicit emotion processing* task assessing the priming (unconscious) effects of emotional expression on participants’ performance of an otherwise neutral face recall task^47^. In the explicit emotion processing task, 48 images from a standardized and validated set of emotional face images ^48^, modified for a consistent luminance, are centrally presented on a computer screen. These images depict 8 individuals (four women and four men) showing either neutral expressions or one of five evoked emotional expressions. Eight images of each expression type are pseudorandomly presented for 2 seconds each, during which participants select, as quickly and accurately as possible, the correct label from a list of emotional expression labels. After an interval of approximately 20 minutes, during which participants complete unrelated cognitive tasks, participants complete the implicit emotion processing task. Stimuli in the implicit task include the 48 face images presented during the explicit task as well as an additional 48 previously unseen images from the same standardized set. Previously seen faces are paired with previously unseen faces matched by sex, emotional expression, and other presentation parameters. From each simultaneously presented image pair, participants are asked to select, as quickly and accurately as possible, the image they have previously seen. As participants are not required to explicitly process or respond to any emotional content, this task assesses the priming effect of the emotional content of faces on this ‘previously seen’ vs. ‘unseen’ detection decision. For both the explicit and implicit emotion processing tasks, Z-scores reflected normed performance on measures of accuracy (% correctly labeled expressions, or % correctly recalled faces) and reaction time. In this analysis our focus was on explicit and implicit processing of expressions of ‘fear’ and ‘happiness’.

### Rodent behavioral tasks

#### Auditory fear conditioning and extinction

All fear conditioning and extinction procedures occurred inside standard operant boxes (Med Associates, St. Albans, VT, USA). On the fear conditioning day, mice were brought into the room individually in white paper boxes, placed into the chambers in Context A, and exposed to 4 tone-shock pairings (30 second, 80 dB, 5 kHz tone; 1 second, 0.5 mA electric shock). Context A consisted of a darkened experimental room, red lights and a slanted plastic ceiling inside the conditioning box, and an electric floor consisting of parallel metal bars. On fear extinction days 2-6, mice were brought into the room in their home cage on a cart, placed into the chambers in Context B, and exposed to 4 tone-only presentations (30 second, 80 dB, 5 kHz tones). Context B consisted of a lit experimental room, yellow lights and a clear flat ceiling inside the conditioning box, a white plastic mat covering the metal grid floor, and Simple Green solution (Huntington Beach, CA, USA) placed in the tray under the metal grid floor. FreezeFrame software (Wilmette, IL, USA) was used to perform tone/shock procedures and to automate quantification of freezing behavior.

#### Elevated zero maze (EZM)

Mice habituated to the behavior room for 30 minutes before behavioral testing. Mice were placed in the middle of the open arm of an EZM in a soundproof enclosure and explored the maze for 15 minutes. Video tracking software (Noldus, Leesburg, VA, USA) was used to quantify time spent in the open and closed arms. EZM data was collected prior to fear conditioning and extinction.

#### Grooming behavior

Mice habituated to the behavior room for 30 minutes before behavioral testing. Mice were placed in a clear cylindrical open field in a soundproof enclosure. Cameras placed above and beside the cylinder were used to collect behavioral data (Noldus) and grooming was manually scored by an experimenter blind to genotype.

#### Goal vs. Habit Lever Press Paradigm

Methods were modified from Gremel et al. 2013 ^49^ to assess the acquisition of instrumental reward learning, goal-directed behavior and habit behavior within a single subject. Mice underwent two training sessions daily in two distinct contexts (clear plastic walls vs. black and white striped walls) in Med Associates operant chambers housed in sound attenuation boxes. The training schedule began with two days of fixed ratio 1-fixed time 30 (FR1-FT30) training (nosepoke for reward or free reward delivered every 30 seconds) followed by continuous reinforcement 15 (CRF15) training (reward contingent upon nosepoke, maximum 15 rewards/60 minutes) until all mice reached the criterion of 15 rewards (BioServ Dustless Precision Pellet F0071, 20 mg, Flemington, NJ, USA) within 60 minutes two days in a row. To prevent overtraining, mice obtaining this criterion sooner were occasionally rested, while those still learning had additional training sessions. Once all mice had reached criterion, mice then ran four days on continuous reinforcement 30 (CRF30) training (reward contingent upon nosepoke, maximum of 30 rewards/60 minutes). Mice underwent FR1-FT30, CRF15, and CRF30 training in each environmental context. This initial training was followed by random ratio (RR) and random interval (RI) training in distinct training contexts that were counterbalanced across mice. Mice underwent one day of random ratio 5 (RR5) training (reward delivered on average every 5 nosepokes with a 0.2 probability that a given nosepoke produces a reward) and random interval 15 (RI15) training (reward delivered upon nosepoke on average every 15 seconds regardless of nosepoke vigor). This was followed by two days of RR10 and RI30 and then four days of RR20 and RI60 training. Mice had 60 minutes to earn a maximum of 15 rewards per training context during the RR/RI training sessions.

### Statistical Analysis

For both explicit and implicit emotion processing tasks, one-sample Wilcoxon rank sum tests (two-tailed, μ=0) were used to test whether the distribution of OCD participants’ age- and gender-normalized z-scores differed from those of the reference population. Additionally, a measure of bias from positive to negative valence was computed by subtracting normed performance for happy faces from normed performance for fearful faces for each measure, and these difference scores were similarly tested via one-sample Wilcoxon rank sum tests (two-tailed, μ=0) to assess whether OCD participants performance differed from the reference population. Lastly, these difference scores were assessed for correlation with clinical measures of OCD (Y-BOCS total, obsessions subscale and compulsions subscale) and with DASS scores, as well as with age and gender using Spearman’s rank correlation coefficient. Spearman partial correlation was used to assess potentially confounding variables. Given the hypothesis-generating nature of the investigation, p-values are reported without correction for multiple comparisons. Analyses were performed using RStudio version 1.2.1335.

Analysis of all rodent data was performed using GraphPad Prism 8. For the fear conditioning and extinction experiment, mice that did not sufficiently learn the tone-shock association as evidenced by an average of less than 40% freezing to the tone on Day 1 of extinction were excluded from the analysis (5 WT and 2 KO mice excluded). For both fear conditioning and operant conditioning analysis, differences in a single numerical variable between SAPAP3 WT and KO mice were assessed using a Wilcoxon rank sum test. Differences in repeated measures of a single numerical variable between genotypes or within individual genotypes were assessed using repeated measures two-way ANOVA. Where appropriate, data were further analyzed using Sidak or Tukey multiple comparisons tests.

## RESULTS

### OCD Patient Sample

N=41 individuals with OCD completed the emotion processing tasks. Demographic and clinical characteristics of the sample are described in Table 1. Average total Y-BOCS score was 26.1 (*s* = 4.7) indicating moderate-to-severe symptoms of OCD. Average depression scale score on the DASS was 13.0 (*s* = 9.8) indicating a mild-to-moderate level of depressive symptoms.

**Table 1:**
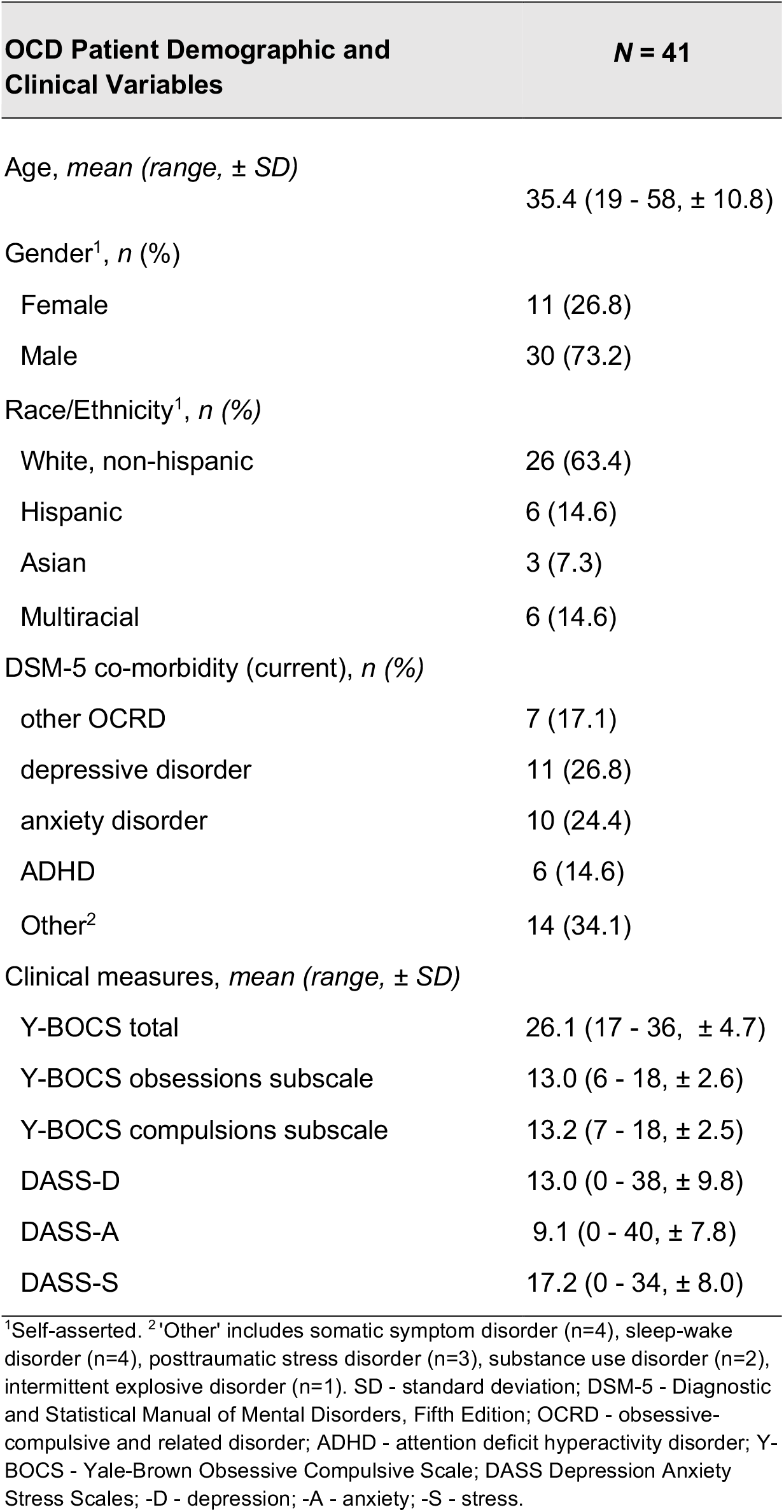

### Explicit Emotion Processing Task

Compared to healthy reference norms, OCD participants were significantly more accurate at identifying fearful expressions and non-significantly less accurate at identifying happy expressions, with a difference score (fearful - happy) representing a bias towards accuracy for fearful faces (**Figure 1A**, Wilcoxon Rank Sum Test, fearful: mean z = 0.47 ±0.12, p = 0.001, happy: mean z = −0.49 ±0.20, p = 0.09, fearful - happy mean z difference = 0.96 ±0.20, p < 0.001). OCD patients were slower in response to both fearful and happy faces than the healthy reference norms, but this difference was statistically significant only for happy faces (**Figure 1B**, Wilcoxon Rank Sum Test, fearful: mean z = −0.31 ±0.20, p = 0.14; happy: mean z = −0.84 ±0.21, p < 0.001). The difference score for explicit identification reaction times (fearful-happy) suggests a relative acceleration of response to fearful versus happy faces (**Figure 1B**, Wilcoxon Rank Sum Test, fearful - happy mean z difference = 0.53 ±0.23, p = 0.025). Overall, these results indicate that on this assessment of explicit, conscious emotion processing, OCD patients respond more effectively to negative versus positive valence stimuli, with both relatively greater accuracy and speed.

**Figure 1.**
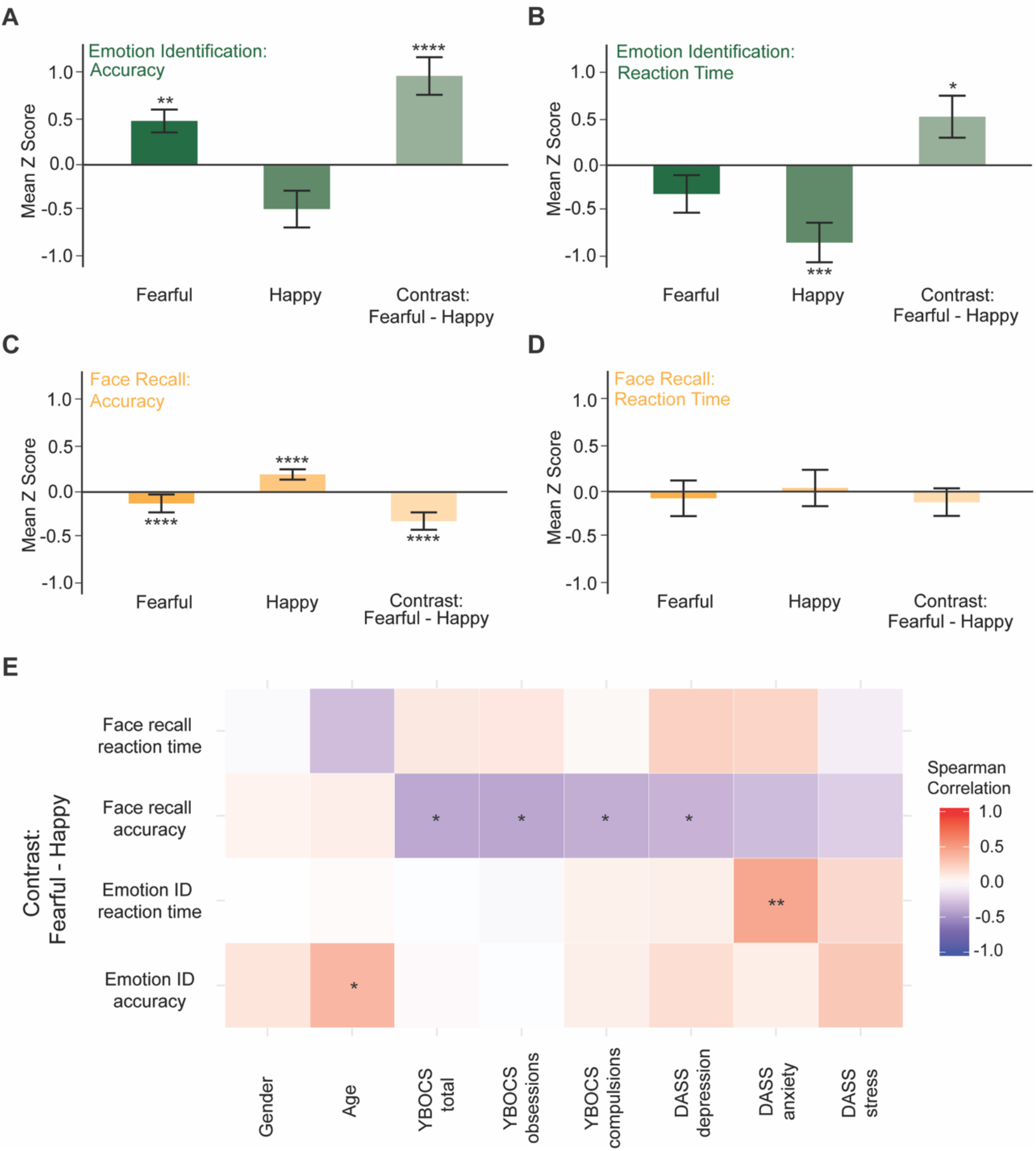
Explicit and implicit emotion processing tasks. *Explicit processing task:* relative to a normative population, (A) OCD patients more accurately identify facial expression of fear, and show bias towards more accurate identification of fearful relative to happy expression. (B) OCD patients more slowly identify facial expression of happiness, and show bias towards more rapid identification of fearful relative to happy expression. *Implicit emotion processing task:* relative to a normative population, (C) OCD patients less accurately recall faces expressing fear, and more accurately recall faces expressing happiness, with negatively biased recall accuracy for faces expressing fear relative to happiness. (D) OCD patients do not differ in speed of face recall, nor is speed of recall biased by the presence of fearful relative to happy expression. (E) Correlations between demographic and clinical variables and fearful vs. happy expression bias on normalized measures of speed and accuracy for both emotion identification (explicit emotion processing) and face recall (implicit emotion processing) tasks. Error bars show standard error of the mean. * = p ≤ 0.05, ** = p ≤ 0.01, *** = p ≤ 0.001, **** = p ≤ 0.0001.

### Implicit Emotion Processing Task

Relative to healthy reference norms, OCD participants were significantly less accurate for face recall when influenced implicitly by fearful expression and more accurate for recall when influenced implicitly by happy expression (**Figure 1C**, Wilcoxon Rank Sum Test, fearful: mean z = −0.12 ±0.10, p < 0.0001, happy: mean z = 0.19 ±0.06, p < 0.0001). The difference score (the contrast of scores for the implicit influence of fearful minus happy faces) suggests that there is an implicit interference of fear on face recall, relative to the implicit facilitation of happiness, with a small to moderate effect size (**Figure 1C**, Wilcoxon Rank Sum Test, fearful - happy mean z difference = −0.32 ±0.10, p < 0.0001). OCD patients did not differ from healthy reference norms in the implicit influence of fearful or happy expression on the reaction time for face recall, nor on the contrast measure of difference for the implicit influence of fearful minus happy expression (**Figure 1D**, Wilcoxon Rank Sum Test, fearful: mean z = −0.07 ±0.20, p = 0.53; happy: mean z = 0.04 ±0.20, p = 0.46, fearful - happy mean z difference = −0.11 ±0.15, p = 0.38). Overall, these results suggest that negative relative to positively valenced stimuli interfere with the performance on a simple recall task in OCD patients even when the influence of these stimuli is implicitly processed.

### Correlations with Clinical and Demographic Variables

To explore whether biases in negative vs. positive valence processing corresponded to clinical phenomena in OCD patients, we assessed correlations between fearful–happy difference measures and demographic and clinical measures. The degree of difference in explicit emotion identification accuracy for fearful vs. happy faces was positively correlated with participant age, though not with any clinical measure (**Figure 1E**, Spearman’s rank correlation coefficient, emotion ID accuracy x participant age: rho = 0.37, p = 0.018). The degree of bias towards faster reaction times to fearful stimuli in the explicit emotion identification task correlated positively with scores for the Anxiety subscale of the DASS, but was not correlated significantly with other clinical or demographic variables (**Figure 1E**, Spearman’s rank correlation coefficient, emotion ID reaction time contrast x DASS anxiety: rho = 0.46, p = 0.003). For the implicit emotion processing task, the bias towards decreased recall accuracy for fearful relative to happy faces correlated with OCD severity by Y-BOCS (including component subscales) as well as with the Depression subscale of the DASS such that greater symptoms on these clinical measures were associated with more potent interference on face recall by fearful relative to happy expression (**Figure 1E**, Spearman’s correlation coefficient, face recall accuracy contrast x Y-BOCS: rho = −0.38, p = 0.015; implicit negative bias x DASS depression: rho = −0.32, p = 0.04). The correlation between negative bias in face recall accuracy and Y-BOCS remained significant even when controlling for DASS depression (rho_partial_= −0.34, p = 0.031). Overall, these results suggest that while subjective anxiety may be associated with faster explicit (conscious) reactivity to negative versus positive valence, clinical OCD symptoms may be associated with greater implicit (non-conscious) impact of negative versus positive valence on other forms of information processing.

### Fear Conditioning and Extinction Task

SAPAP3 KO mice showed significantly increased grooming behavior (**Figure 2A**, Mann-Whitney U Test, p = 0.0007; WT = 12, KO = 11) as well as higher innate anxiety (**Figure 2B**, Mann-Whitney U Test, p = 0.0426; WT = 6, KO = 8) compared to WT controls, replicating established phenotypes and confirming the validity of the model. We next assessed alterations in negative valence processing in SAPAP3 KO mice using a classical auditory fear conditioning paradigm as well as a multi-day fear extinction procedure (**Figure 2C**). During fear conditioning, SAPAP3 KO mice showed no differences in freezing to tone 1, but showed a significant increase in freezing to tones 2-4 compared to WT mice (**Figure 2D**, Two-way Repeated Measures ANOVA, Time x Genotype p < 0.05, Time p < 0.0001, Genotype p = 0.0001; Sidak’s Multiple Comparisons Test, WT vs. KO Tone 1 p = 0.9893, Tone 2 p < 0.05, Tone 3-4 p ≤ 0.001; WT = 11, KO = 11). These results demonstrate that SAPAP3 KO mice show enhanced fear learning.

**Figure 2.**
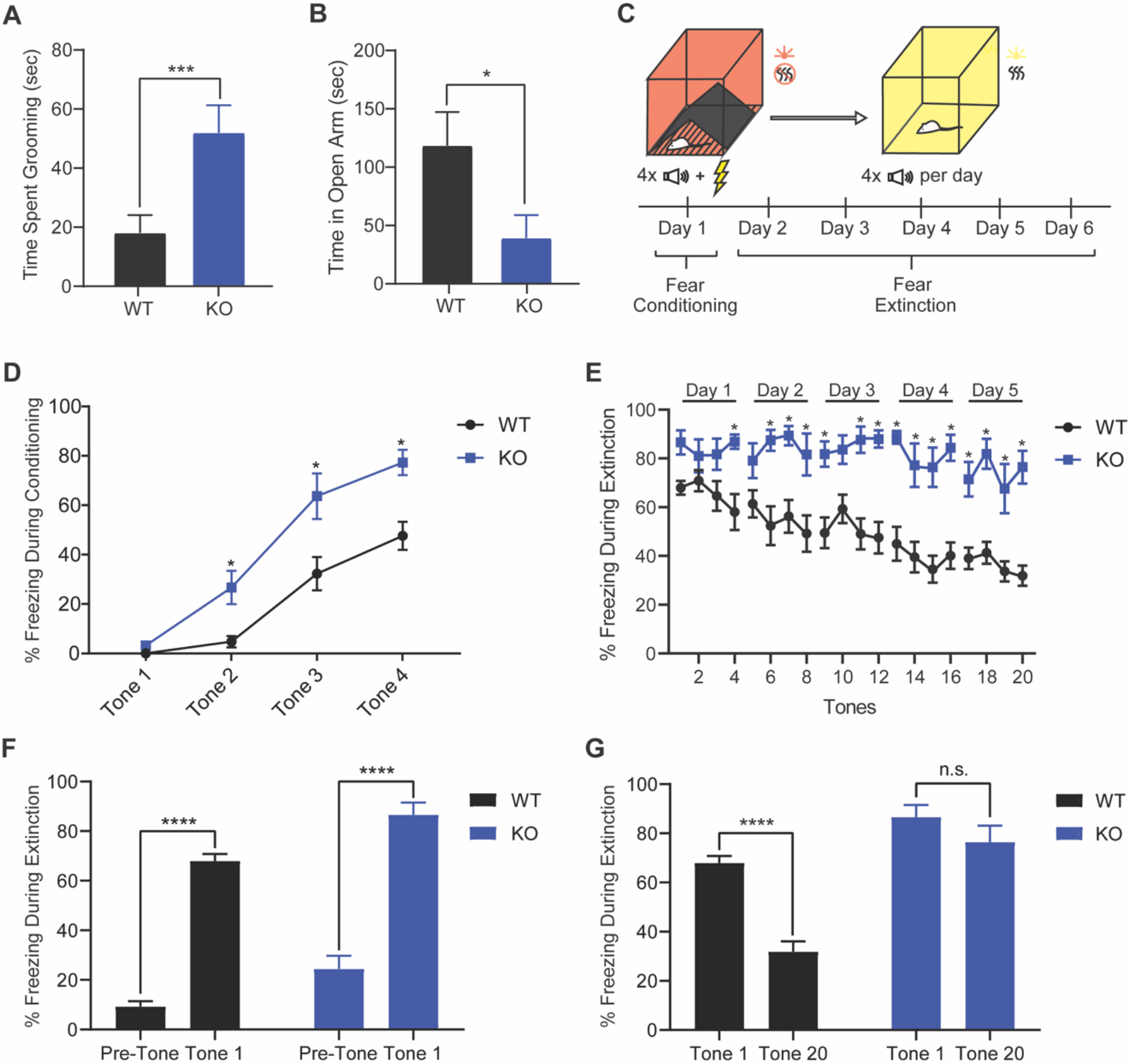
SAPAP3 KO mice show enhanced fear learning and impaired fear extinction. (A) SAPAP3 KO mice spent significantly more time grooming than WT mice. (B) SAPAP3 KO mice spent less time in the open arm of the elevated zero maze (EZM) than WT mice. (C) Schematic for the fear conditioning and extinction procedure. (D) SAPAP3 KO mice froze to the tone significantly more than WT mice during tones 2-4 of fear conditioning. (E) SAPAP3 KO mice froze to the tone significantly more than WT mice during days 2-5 of fear extinction. (F) Both SAPAP3 WT and KO mice froze significantly more during the first tone presentation than during the pre-tone period on the first day of extinction. (G) SAPAP3 WT mice significantly decreased their freezing from the first to the last tone presentation during extinction while SAPAP3 KO mice did not show a significant decrement in freezing across extinction; * = p ≤ 0.05, ** = p ≤ 0.01, *** = p ≤ 0.001, **** = p ≤ 0.0001; stars in graphs 2D and 2E convey significance but not level of significance.

During extinction, following the first few tones, WT mice showed significantly lower freezing to the tone compared to SAPAP3 KO mice (**Figure 2E**, Two-way Repeated Measures ANOVA, Time × Genotype p < 0.01, Time p < 0.0001, Genotype p < 0.0001; Sidak’s Multiple Comparisons Test, WT vs. KO Tone 4 p < 0.05, Tone 6-9 p < 0.01, Tone 11 p < 0.001, Tone 12-13 p < 0.0001, Tone 14 p < 0.001, Tone 15-16 p < 0.0001, Tone 17 p < 0.01, Tone 18 p < 0.0001, Tone 19 p < 0.01, Tone 20 p < 0.0001; WT = 11, KO = 11). Importantly, both WT and SAPAP3 KO mice successfully recalled the tone-shock association on Day 1 of extinction, as evidenced by significantly higher freezing to the tone than freezing during the pre-tone period (**Figure 2F**, Two-way Repeated Measures ANOVA, Time x Genotype p = 0.6648, Time p < 0.0001, Genotype p < 0.001; Sidak Multiple Comparisons Test, WT Pre-Tone vs. Tone 1 p < 0.0001, KO Pre-Tone vs. Tone 1 p < 0.0001; WT = 11, KO = 11). WT mice significantly decreased their freezing to the tone from early to late extinction, which indicates successful fear extinction. Conversely, SAPAP3 KO mice did not show a significant difference in freezing between early and late extinction, suggesting impaired ability to extinguish fear (**Figure 2G**, Two-way Repeated Measures ANOVA, Time x Genotype p < 0.05, Time p < 0.0001, Genotype p < 0.0001; Sidak’s Multiple Comparisons Test, WT Tone 1 vs. Tone 20 p < 0.0001, KO Tone 1 vs. Tone 20 p = 0.2612; WT = 11, KO = 11).

To better characterize the nature of altered fear processing in the SAPAP3 KO mice, we plotted freezing behavior in WT and SAPAP3 KO mice across the entire trial session for extinction day 2 (**Figure 3A**), which is when differences in cued freezing emerged between WT and SAPAP3 KO mice. SAPAP3 KO mice also showed greater non-cued freezing throughout the entire trial period (**Figure 3A**). Quantifying differences in freezing behavior across all extinction days, we found that SAPAP3 KO mice showed significantly increased freezing during the pre-tone period (**Figure 3B**, Mann-Whitney U Test, p = 0.0041; WT = 11, KO = 11) and during the period in between tones (**Figure 3C**, Mann-Whitney U Test, p = 0.0003; WT = 11, KO = 11) compared to WT mice. This general increase in freezing behavior was due to an increase in freezing bout duration (**Figure 3D**, Mann-Whitney U Test, p = 0.001; WT = 11, KO = 11) with no difference in the total number of freezing bouts (**Figure 3E**, Mann-Whitney U Test, p = 0.7348; WT = 11, KO = 11). Overall, these results indicate that SAPAP3 KO mice show elevations in fear processing beyond cued fear.

**Figure 3.**
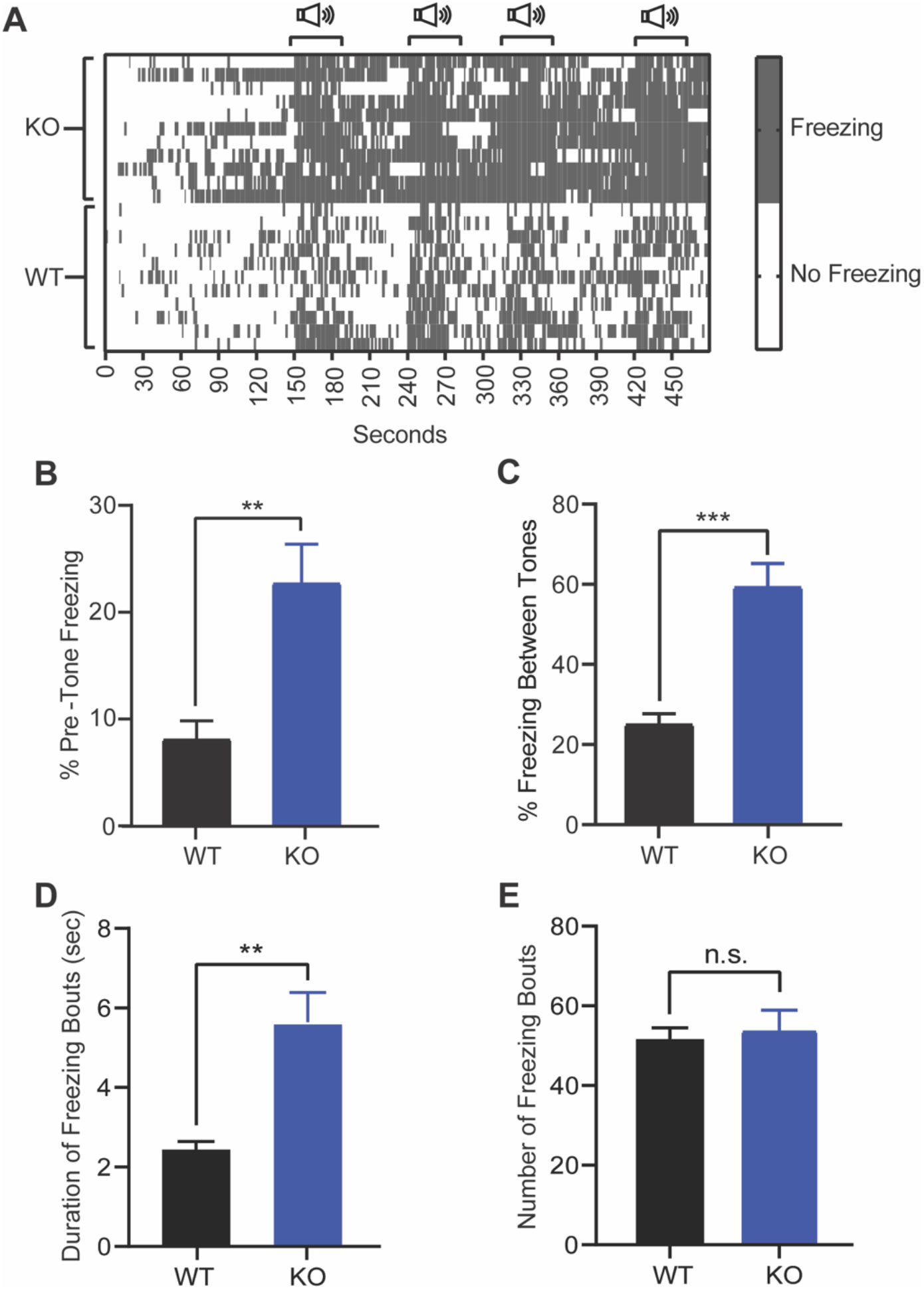
SAPAP3 KO mice show alterations in fear processing beyond cued fear. (A) Heatmap showing all SAPAP3 WT and KO freezing events across the entire trial session for extinction day 2. Individual freezing events within one second time bins are demarcated with a grey dash. Each row represents the freezing bouts of an individual animal. (B) SAPAP3 KO mice freeze significantly more during the pre-tone period than WT mice throughout extinction. (C) SAPAP3 KO mice spent significantly more time freezing between tones than WT mice throughout extinction. (D) SAPAP3 KO mice had a significantly increased freezing bout duration compared to WT mice. (E) SAPAP3 KO mice showed no difference in freezing bout number compared to WT mice; n.s. = p > 0.05, * = p ≤ 0.05, ** = p ≤ 0.01, *** = p ≤ 0.001, **** = p ≤ 0.0001.

### Reward-Based Operant Conditioning Task

To assess alterations in positive valence processing, we trained WT and SAPAP3 KO mice on a multi-stage reward-based operant conditioning paradigm (**Figure 4A**). WT mice learned the association between nosepoke and reward quickly and then stably maintained nosepoke behavior, whereas SAPAP3 KO mice were much slower at learning nose-poke behavior, with two KO mice never achieving stable performance (**Figure 4B**, Two-way Repeated Measures ANOVA, Genotype x Training Day p < 0.0001, Training Day p < 0.0001, Genotype p < 0.0001; Sidak’s multiple comparisons, WT vs. KO Day 2 p < 0.001, for Days 3-12 p < 0.0001, Day 14 p < 0.01, Day 15 p < 0.001, Day 17 p < 0.05; WT = 11, KO = 8). It took SAPAP3 KO mice on average over 3 times as many training days to establish stable nosepoke behavior compared to WT mice (**Figure 4C**, Wilcoxon Rank-Sum Test, p < 0.001; WT = 11, KO = 6). After examining differences in acquisition of nosepoke behavior, we next examined how nosepoke behavior was maintained in each genotype under two different reinforcement schedules, one biasing for habit formation (random interval), and one for goal-directed behavior (random ratio) ^49^. SAPAP3 KO mice maintained their learned nosepoke behavior under the reinforcement schedule that promotes habitual responding, but their ability to earn reward greatly diminished under the reinforcement schedule that promotes goal-directed responding (**Figure 4D**, Two-way Repeated Measures ANOVA, Genotype × Training Day p < 0.0001, Training Day p < 0.0001, Genotype p < 0.0001; Tukey’s multiple comparisons, day 1 of RR10/RI30 training WT RI vs. KO RR p < 0.001 and WT RR vs. KO RR p < 0.05, day 2 of RR10/RI30 training WT RI vs. KO RR p < 0.001, WT RR vs. KO RR p < 0.01, and KO RI vs. KO RR p < 0.01, for each RR20/RI60 training days 1 through 4 WT RI vs. KO RR p < 0.0001, WT RR vs. KO RR p < 0.0001, and KO RI vs. KO RR p < 0.0001; WT = 11, KO = 6). These data indicate that SAPAP3 KO mice have a selective learning deficit under positive reward conditions with difficulties both in acquiring positive-reinforced behaviors and in maintaining goal-directed responding for rewards.

**Figure 4.**
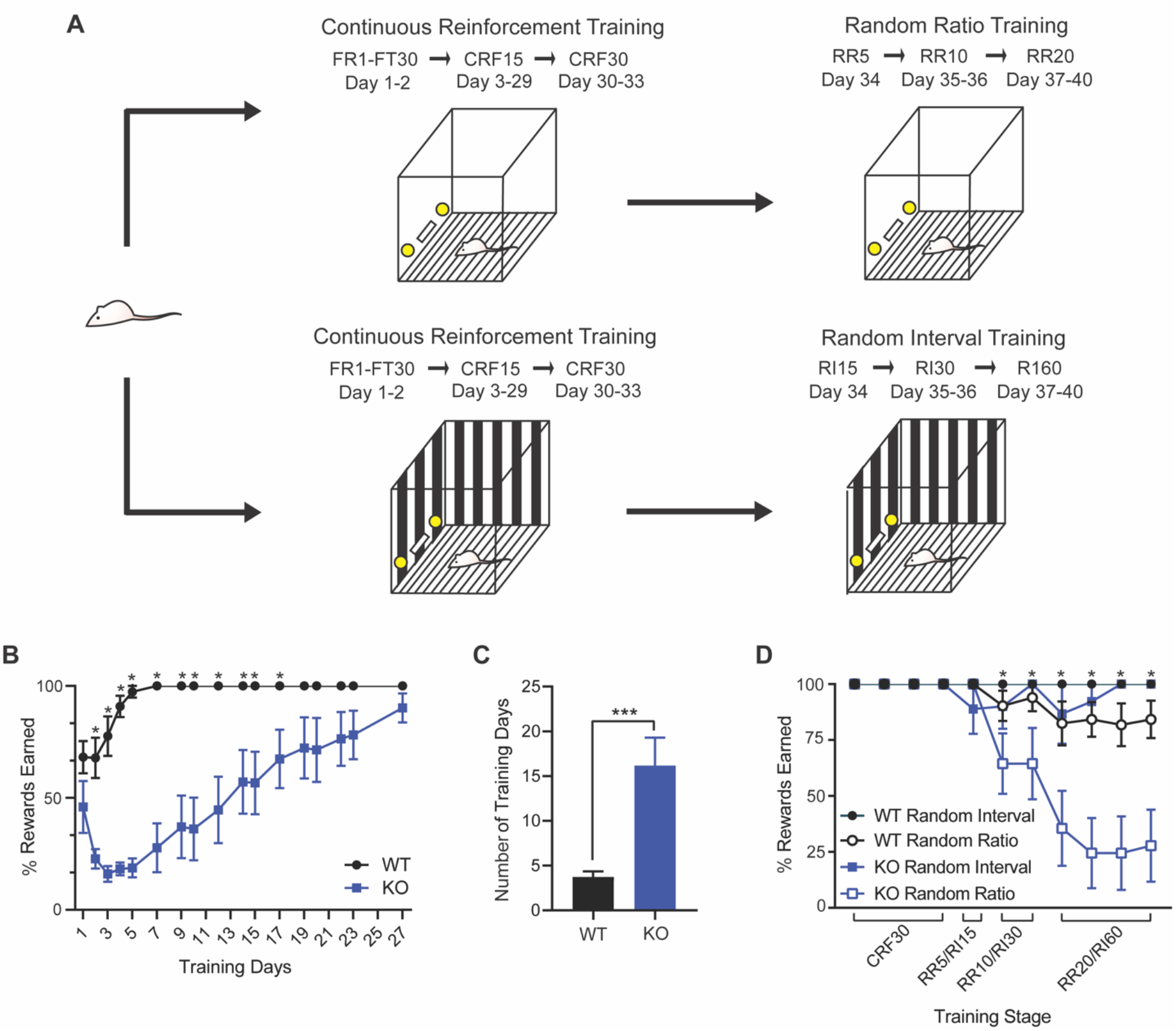
SAPAP3 KO mice show impaired acquisition of reward learning and goal-directed behavior. (A) Schematic for operant conditioning procedure. (B) Learning curve for the acquisition of CRF15 nosepoke behavior on days for which both WT and KO mice received training time. (C) KO mice took significantly longer than WT mice to achieve stable nosepoke behavior for reward during CRF15 training. (D) KO mice struggled engaging in goal-directed nosepoke reward schedule (RR), but did not have difficulty earning rewards under a habit-promoting nosepoke reward schedule (RI); FR, fixed ratio; FT, fixed time; CRF, continuous reinforcement; RR, random ratio; RI, random interval; * = p < 0.05, ** = p < 0.01, *** = p < 0.001, **** = p < 0.0001; stars in graphs 4B and 4D convey significance but not level of significance.

## DISCUSSION

We uncovered key alterations in positive and negative valence processing in OCD patients and in a preclinical rodent model of compulsive behavior, the SAPAP3 KO mouse model, showing promise for translation across species. On behavioral measures, OCD patients were found to have speeded and more accurate identification of fearful compared to happy facial expressions and a greater implicit influence of fear versus happiness on an otherwise neutral face recall task. Speeded explicit responses to fear were correlated with greater subjective anxiety while greater implicit influence of fear was correlated with severity of OCD symptoms. In the rodent model, we found that SAPAP3 KO mice showed enhanced fear learning, impaired fear extinction, impaired acquisition of reward learning, and impaired goal-directed behavior. Overall, our results suggest that OCD patients and the SAPAP3 KO mouse model show parallel trends toward heightened negative valence processing and impaired positive valence processing, suggesting the translational validity of our valence processing assays.

The profile of response to fearful versus happy expressions in OCD patients is indicative of a negative valence bias. Previous studies of facial emotion processing in OCD have largely focused on whether OCD patients have deficits in identifying symptom-congruent expressions of disgust and have yielded mixed results^50–54^. These studies have not generally explored valence effects, however, and have typically assessed patients taking psychotropic medication, whereas antidepressant medication has been shown to alter processing of emotional expressions in patients with OCD ^55^ and with depression ^56^. Established assessments of negative valence processing, specifically tests of attentional bias such as the emotional stroop or dot-probe task, characteristically show attentional bias towards threat stimuli in anxiety disorders, yet have produced equivocal findings in OCD ^57–59^. The inclusion of both explicit and implicit emotion processing measures is a strength of our study, and our observation that anxiety symptoms and OCD symptoms show distinct patterns of correlation with explicit versus implicit measures may help clarify potential differences in negative valence processing between OCD and anxiety disorders.

Our observation that the implicit influence of fear is correlated with OCD symptom severity should be interpreted cautiously given the potential for Type I error, yet this correlation nonetheless suggests that measures of implicit emotion processing may be more sensitive to OCD phenomena than those of explicit emotion processing. In our study, for OCD patients, resources devoted to unconscious processing of negative valence stimuli may have detracted from the consciously-attended non-emotional recall task, leading to relative performance deficits in the non-emotional task. This model is supported by replicated neuroimaging findings that OCD patients show greater activation of the amygdala and other regions involved in a face matching task similarly assessing the implicit influence of fear versus happiness, with the degree of activation correlated with severity of obsessive-compulsive symptoms per Y-BOCS ^60,61^. Alternatively, a potential correlation between OCD symptoms and poorer recall accuracy for fearful faces may be related to the increased generalization of fear stimuli seen in individuals with obsessive-compulsive traits ^62^, in line with evidence in OCD patients of decreased engagement of hippocampus during fear conditioning ^9^. Although OCD patients have been shown to have deficits in implicit learning generally ^63^, the differences in recall accuracy for fearful vs happy faces observed in our study argues against a mechanism independent of emotional valence.

The correlation between speeded responses to fear and severity of anxiety is in line with findings suggesting that anxiety biases responses to affect-congruent stimuli. This association has been shown via increased accuracy of identification of fearful faces by individuals with high trait anxiety ^64,65^ (although other studies have failed to replicate this finding ^66,67^), and by increased accuracy of fearful face identification by healthy volunteers under threat of electrical shock, a condition that otherwise degrades accuracy of identification of neutral or happy faces ^68^. A specific speeding of reaction time for identifying fearful faces has also been observed in adolescents with depressive and anxiety symptoms ^69^.

In parallel to our clinical findings, the SAPAP3 KO mice also show enhanced sensitivity to negative valence. SAPAP3 KO mice displayed both enhanced fear learning and impaired fear extinction, though only impairments in fear extinction have been consistently found in OCD patients ^9–12^. This suggests that future studies should focus on the fear extinction phenotype in SAPAP3 KO mice to increase translational potential. SAPAP3 KO mice also showed impairments in the acquisition of reward learning which have similarly been reported in OCD patients ^18,19,21^. However, these acquisition abnormalities have been found in some but not all studies ^22,25^, which may be due to differences in medication status ^21^. Finally, our study revealed that SAPAP3 KO mice have impaired goal-directed learning and intact habit learning. Studies have shown that OCD patients not only show impaired goal-directed behavior, but also excessive habit formation as evidenced by improper responding after outcome devaluation ^22–25^.

While alterations in cued fear, acquisition of reward learning, and goal-directed learning all mimic clear phenomena seen in OCD patients, the clinical relevance of the enhanced non-cued fear seen in the SAPAP3 KO mice is less clear. The elevated pre-tone freezing we observed in SAPAP3 KO mice on extinction days could indicate fear generalization to the novel ‘safe’ context, which could mimic OCD patients’ inability to discriminate between safe and threatening stimuli ^6,70^. Additionally, the elevated freezing between tones seen in the SAPAP3 KO mice could be evidence that SAPAP3 KO mice have an underlying propensity to get ‘stuck’ in certain behaviors similar to OCD patients’ difficulty disengaging from repetitive thoughts and actions. The underlying structure of the freezing bouts in the SAPAP3 KO mice, which show longer freezing bout lengths with no change in freezing bout number, support this hypothesis.

It is important to note that differences in task design may limit the parallels that can be drawn across tasks. While our negative valence paradigm involved fear conditioning which relies on associative learning, our positive valence paradigm involved a reward-based operant conditioning task which relies on both associative and instrumental learning. In addition, the clinical study is distinct from the animal tasks in that the rodent-based assays involved learning associations between conditioned stimuli and rewarding or aversive outcomes while the clinical study involved explicit and implicit emotion processing. Future studies should develop tasks that incorporate both valences and are as similar as possible in rodents and humans to increase translational potential.

We have shown that OCD patients and the SAPAP3 KO mouse model of compulsive behavior have consistent trends toward enhanced negative valence processing and impaired positive valence processing. We also find that valence assay results seen in OCD patients correlate with metrics of OCD symptom severity. Overall, our results highlight the importance of tracking valence processing alterations in OCD patients in clinical settings and confirm the clinical relevance of studying valence processing abnormalities in the SAPAP3 mouse model. Future work building upon our behavioral foundation has the potential to greatly aid in refining current therapies and identifying novel treatment strategies for OCD patients.

## Conflict of Interest

In the last 3 years, Dr. Rodriguez has served as a consultant for Epiodyne and received research grant support from Biohaven Pharmaceuticals and a stipend from APA Publishing for her role as Deputy Editor at The American Journal of Psychiatry. Dr. Williams has served as a scientific advisor for One Mind Psyberguide, a member of the executive advisory board for the Laureate Institute for Brain Research and holds patent 16921388 (Systems and Methods for Detecting Complex Networks in MRI Image Data) unrelated to the present study. All other authors declare that they have no conflicts of interest.

## Funding

Preparation of this work was supported in part by IOCDF Breakthrough Award and NIMH (R01MH105461) to Dr. Rodriguez, and a Chan Zuckerberg Biohub Investigator Award to Dr. Gunaydin. Dr. van Roessel is supported by the Office of Academic Affiliations, Advanced Fellowship Program in Mental Illness Research and Treatment, Department of Veterans Affairs, and by a Miller Foundation Award for Psychiatric Research.

